# TMT-based proteomic analysis of the antibiotic effects of ShangKeHuangShui against *Straphylococcus aureus*

**DOI:** 10.1101/2021.07.22.453437

**Authors:** Lichu Liu, Na Zhao, Kuangyang Yang, Honghong Liao, Xiaofang Liu, Ying Wu, Lei Lei, Xiao Peng, Yuanyan Wu

## Abstract

Post-traumatic infection is a serious orthopedic trauma complication commonly caused by *Staphylococcus aureus* (SAU). ShangKeHuangShui, also known as Yellow Aqueous Concentrate of Traumatology Herbs (YACTH), is prepared from six Chinese herbal medicines, which has been used for decades in our hospital to prevent post-traumatic infection.

In the present study, we investigated the *in vitro* antibacterial effects and underlying mechanism of YACTH against SAU. YACTH exhibited significant antibacterial activity against SAU with the minimum inhibitory concentration (MIC) of 8.625 mg/mL. Proteomic analysis based on Tandem mass tag (TMT) showed different protein expression levels in SAU under the YACTH and control conditions. Compared to the control group, the expression level of 490 proteins of YACTH treated group significantly changed (>1.2 fold, P <0.05). Biological informatics analysis showed that these differential proteins were widely involved in a variety of biological processes and multiple metabolic pathways. We then selected 26 target proteins to further conduct Parallel reaction monitoring (PRM). The results revealed that the 9 down-regulated proteins were mainly involved in RNA polymerase, oxidative phosphorylation, ABC transporter and glycolysis; whereas, the 17 up-regulated proteins are related to RNA degradation, mismatch repair and ribosome.

In conclusion, YACTH, which contains the water-soluble components of six Chinese herbs, is first reported to exhibit antibacterial activity against SAU *in vitro*. The results provide novel insights into the antibacterial mechanisms of YACTH on protein networks, which may help us find new potential antibiotic targets.

## Introduction

*Staphylococcus aureus* (SAU), one species of coagulase Gram-positive staphylococcus, is widely found in the skin of mammals [1]. Due to its infectiousness and resistance to high temperature [2], SAU has become a major nosocomial pathogenic cause of superficial lesions, systemic infections, and several toxemic syndromes [3]. Skin injuries with open wounds can be easily infected by SAU and result in impetigo and folliculitis [4]. Infection with SAU is a serious complication for orthopedics trauma and is quite common in open fracture management [5].

To date, a variety of antibiotics have been invented to fight against SAU infection. Yellow Aqueous Concentrate of Traumatology Herbs (YACTH) is developed by six Chinese herbal medicines including Coptis Chinensis, Philodendron Chinensis, Gardenia jasminoides, Lithospermum, peppermint, and alum [6]. Its water-soluble components of berberine, geniposide, palmatine, and Phellodendron are potential natural antibacterial compounds that are reported to reduce subside swelling, invigorate blood circulation, promote tissue regeneration and improve wound healing [7,8]. However, whether such improvement was achieved by anti-inflammatory or antibacterial effects remained unknown, as well as the relevant molecular mechanism. In the presence of microorganisms, soft tissue injury and the consequent decline of antibacterial capability are the main causes of infection [9]. Considering YACTH contains antibacterial components, suggested that YACTH may promote wound healing through suppressing bacterial activities. Typically, the single component chemical compound, such as beta-lactam antibiotics inhibit SAU through bifunctional transglycosylase-transpeptidases PBP [10] and tetracyclines that bind to the 30S subunit of the ribosome to interfere with the protein synthesis of SAU [11], solely have one molecular target. Previous studies provided many antibiotics with clear mechanisms. The majority of those antibiotics suppress SAU through targeting the cell envelope and proteins that are involved in protein synthesis and nucleic acids biosynthesis [12].

YACTH, as a compound formulation containing numbers of active antibacterial components, might have multi-target antibacterial activities, which makes it difficult to study the mechanism of how YACTH inhibits SAU. Proteomics is the analysis of the entire protein complement of a cell, tissue, or organism under a specific, defined set of conditions [13]. Up to date, proteomics could be applied to many fields due to the rapid development of technologies including protein fractionation techniques, mass spectrometry (MS) technology, bioinformatics, etc. The protein expression changes could be detected through MS, and reveal pathway information through informatics [14].

We hypothesized that YACTH inhibits SAU by regulating a variety of biological processes and multiple metabolic pathways. In the present study, we employed proteomics analysis to provide novel insights into the antibacterial mechanisms of YACTH on protein networks and explore some potential antibiotic targets.

## Materials and methods

### Evaluation of Antibacterial Activity of YACTH

#### Strain and Culture Condition

The strain *S*.*aureus* ATCC 25923 used in this study was kept in lab and cultured aerobically in broth medium(Qingdao Hope Bio-Technology Co., Ltd)on a shaking incubator at 250 rpm, 37 °C.

#### Antibacterial Activity of YACTH

Antibacterial activity of YACTH was tested according to the agar diffusion method as reported previously with slight modification [15]. The YACTH extracts were filtered through 0.22 µm sterilizing filters. Briefly, 100 µL bacterial suspension containing 10^6^ CFU/mL of SAU was evenly smeared on Mueller-Hinton ager plates. Oxford cups (outside diameter of about 8.0 mm) were placed on the plates.After 20 min, 80 µL of YACTH were added to the Oxford cups. Saline was used in parallel experiments as a negative control. The diameters of the inhibitory zones (DIZs) were measured after incubation of the plates at 37 °C for 24 h, which were expressed in millimeter (mm) and recorded as mean ± standard deviation (SD). DIZ values less than 8 mm were considered as no inhibition zone (NIZ). All experiments were performed in triplicate.

#### Determination of MIC and MBC

Minimum inhibitory concentration (MIC) and minimum bactericide concentration (MBC) were determined by a micro-dilution test according to the method described by Jeong [16]. Studies were conducted according to the guidelines recommended by the Clinical and Laboratory Standards Institute (CLSI) [17]. Two-fold serial dilutions of YACTH were prepared in sterile broth media ranging from 0.54-34.5 mg/mL. Afterwards, 180 µL of diluted YACTH was mixed with 20 µL bacterial suspension (approximately 10^7^ CFU/mL) in the 96-well plates. The negative control contained 180 µL diluted YACTH with 20 µL 0.9% NaCl. The plates were incubated at 37 °C for 24 h. The MIC was defined as the concentration at which the corresponding well showed no visible bacterial growth after incubation at 37 °C for 24 h. Afterwards, a subculture of 50 µL from each well with no visible bacterial growth was directly incubated onto the nutrient agar plate incubating at 37 °C for 24 h, and the lowest on centration at which there was no colony growth was defined as MBC.

### Biochemistry experiments

#### Protein extraction and digestion

The proteins were lysed and extracted with SDT buffer (4% SDS,100 mM Tris-HCl,1 mM DTT, pH 7.6), and then quantified with the BCA protein assay kit (Bio-Rad, USA). After trypsin digestion, the peptides of each sample were desalted and concentrated by vacuum centrifugation and reconstituted in formic acid (40,0.1% (v/v)).

#### Filter-aided sample preparation (FASP Digestion)

200 μg of proteins of each sample were treated with 30 μL SDT buffer (4% SDS, 100 mM DTT, 150 mM Tris-HCl pH 8.0). Used UA buffer (8 M Urea, 150 mM Tris-HCl pH 8.0) to remove detergent, DTT, and other low-molecular-weight components by repeated ultrafiltration (Microcon units, 10 kD). Added 100 μL iodoacetamide (100 mM IAA in UA buffer) to block reduced cysteine residues. Incubated the samples for 30 min in a dark environment. Washed the filters with 100 μL UA buffer three times and then 100 μL 25mM NH_4_HCO_3_ buffer twice. Digested protein suspensions with 4 μg trypsin (Promega) in 40 μL 25 mM NH_4_HCO_3_ buffer at 37 °C overnight. The resulting peptides were collected as a filtrate. The peptides of each sample were desalted on C18 Cartridges (Empore™ SPE Cartridges C18 (standard density), bed I.D. 7 mm, volume 3 mL, Sigma), concentrated by vacuum centrifugation and reconstituted in 40 µL of 0.1% (v/v) formic acid. Estimated the peptide content by UV light spectral density at 280 nm(extinctions coefficient= 1.1, 0.1% (g/L)).

#### SDS-PAGE

Mixed 20 µg of protein for each sample with loading buffer (5X) and boiled it for 5 min. The proteins were separated on 12.5% SDS-PAGE gel (constant current 14 mA, 90 min), and stained by Coomassie Blue R-250.

#### TMT Labeling

Peptides (100 μg) of each sample were labeled with TMT reagent according to the manufacturer’s instructions (Thermo Scientific).

#### High pH Reversed-Phase Fractionation

Labeled peptides were fractionated by High pH Reversed-Phase Peptide Fractionation Kit (Thermo Scientific). The dried peptides were reconstituted and acidified with 0.1% TFA solution and loaded on the equilibrated, high-pH, reversed-phase fractionation spin column. Peptides were attached to the hydrophobic resin under aqueous conditions and desalted by water through low-speed centrifugation. Collected 10 fractions from the gradient elution of the attached peptides by increasing acetonitrile concentrations under high-pH. Desalted the collected fractions on C18 Cartridges (Empore™ SPE Cartridges C18 (standard density), bed I.D. 7 mm, volume 3 mL, Sigma) and concentrated them by vacuum centrifugation.

#### LC-MS/MS analysis

Performed LC-MS/MS analysis on a Q Exactive mass spectrometer (Thermo Scientific) that was coupled to Easy nLC (Proxeon Biosystems, now Thermo Fisher Scientific) for 60 min. At a flow rate of 300 nL/min, the peptides were applied to a reverse phase trap column (Thermo Scientific Acclaim PepMap100, 100 μm*2 cm, nanoViper C18) connected to the C18-reversed phase analytical column (Thermo Scientific Easy Column, 10 cm long, 75 μm inner diameter, 3 μm resin) in buffer A (0.1% Formic acid) and separated with a linear gradient of buffer B (84% acetonitrile and 0.1% Formic acid). The mass spectrometer was operated in positive ion mode. MS data were acquired via the most abundant precursor ions from the survey scan (300–1800 m/z) for HCD fragmentation using a data-dependent top10 method dynamically. Set automatic gain control (AGC) target to 3e6 and maximum inject time to 10 ms with a dynamic exclusion duration of 40.0 s. Acquired survey scans at a resolution of 70,000 at m/z 200 and set resolution for HCD spectra to 17,500 at m/z 200, and isolation width was 2 m/z. Normalized collision energy was 30 eV and the underfill ratio, which specifies the minimum percentage of the target value likely to be reached at maximum fill time, was defined as 0.1%. Ran the instrument with peptide recognition mode enabled.

#### Identification and quantitation of proteins

Searcher the MS raw data for each sample using the MASCOT engine, Uniprot (Matrix Science, London, UK; version 2.2) embedded into Proteome Discoverer 1.4 software for identification and quantitation analysis.

## Bioinformatic analysis

### Cluster analysis

Performed hierarchical clustering analysis through Cluster 3.0 (http://bonsai.hgc.jp/~mdehoon/software/cluster/software.htm) and Java Treeview software (http://jtreeview.sourceforge.net) and selected Euclidean distance algorithm for similarity measure and average linkage clustering algorithm (clustering uses the centroids of the observations) for clustering.

### Subcellular localization

A multi-class SVM classification system, CELLO (http://cello.life.nctu.edu.tw/), was used to predict protein subcellular localization.

### Domain annotation

Search protein sequences via the InterProScan software. Identified protein domain signatures from the InterPro member database P fam.

### GO annotation

Locally searched the protein sequences of the selected differentially expressed proteins using the NCBI BLAST+client software (NCBI-blast-2.2.28 ± win32.exe) and found homologue sequences on InterProScan. Mapped then gene ontology (GO) terms and annotated the sequences using the software program Blast2GO. Plotted the GO annotation results by R scripts.

### KEGG annotation

Blasted the studied proteins against the online Kyoto Encyclopedia of Genes and Genomes (KEGG) database (http://geneontology.org/) to retrieve their KEGG orthology identifications. Subsequently mapped the matched proteins to pathways in KEGG.

### Enrichment analysis

The whole quantified proteins were considered as background dataset. The enrichment analysis was applied based on Fisher’s exact test. Further applied Benjamini-Hochberg correction for multiple testing to adjust derived p-values. Only considered functional categories and pathways with p-values under a threshold of 0.05 as significant.

### Statistical Analysis

Statistical analysis was conducted with two-tailed t-test. The P value was set at < 0.05. Fisher’s exact test was used to calculate the P value compare the distribution of each GO classification or KEGG pathway in the target protein set and the total protein set, and the enrichment analysis of GO annotation or KEGG pathway annotation was performed on the target protein set.

## Results

### Antibacterial Activity of YACTH

The antibacterial activity of YACTH against SAU was measured by Oxford cup and micro-dilution methods, diameters of inhibition zone (DIZ) were appreciated as follows: not sensitive (diameter ≤ 8.0 mm), moderately sensitive (8.0 < diameter ≤ 14.0 mm), sensitive (14.0 < diameter ≤ 20.0 mm), and extremely sensitive (diameter >20.0 mm) [15]. The mean of DIZ values was 28.04mm (± 0.94 mm), and the corresponding MIC and MBC against SAU were 8.625 mg/mL and 17.25 mg/mL, respectively.

### LC-MS/MS Analysis

In this study, the TMT technique was applied to analyze the proteomics of SAU treated with YACTH. The experimental procedure is illustrated in Fig 1.

**Fig 1.**
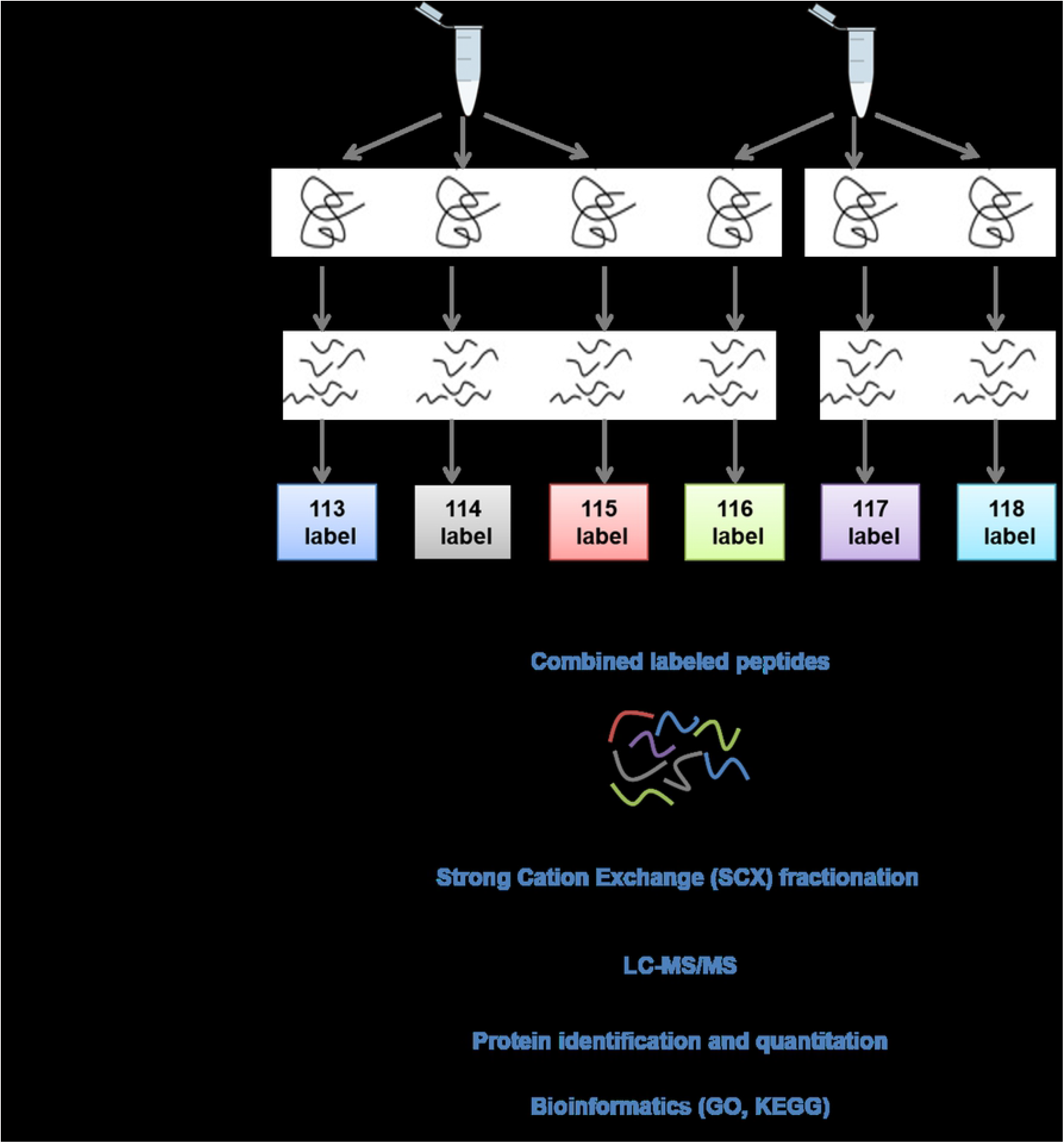
Experimental procedure of LC-MS/MS analysis.

Fig 1 shows the experimental procedures of our study. The YACTH group was compared to the control group. Each group contained three samples and went through protein preparation, labeling, fractionation, LC-MS/MS, protein identification and quantification, and bioinformatic analysis.

In LC-MS/MS analysis, a total of 470327 products were detected on the spectrum with 63749 matched with recorded data. Among these fractionated products, a total of 23544 peptides were identified and 16490 were recognized as unique, while 2583 proteins were quantified out of the 2605 identified proteins. Compared to the control group, the expression of 490 proteins was significantly affected by the treatment of YACTH (P < 0.05), while other proteins remained to be unchanged. Among these proteins, 241 proteins were up-regulated and 249 proteins were down-regulated (Fig 2A). To confirm the regulatory effects of YACTH on the growth and metabolism of SAU, a cluster plot was constructed to demonstrate the significant difference in protein expression between the control group and the YACTH treatment group (Fig 2B).

**Fig 2.**
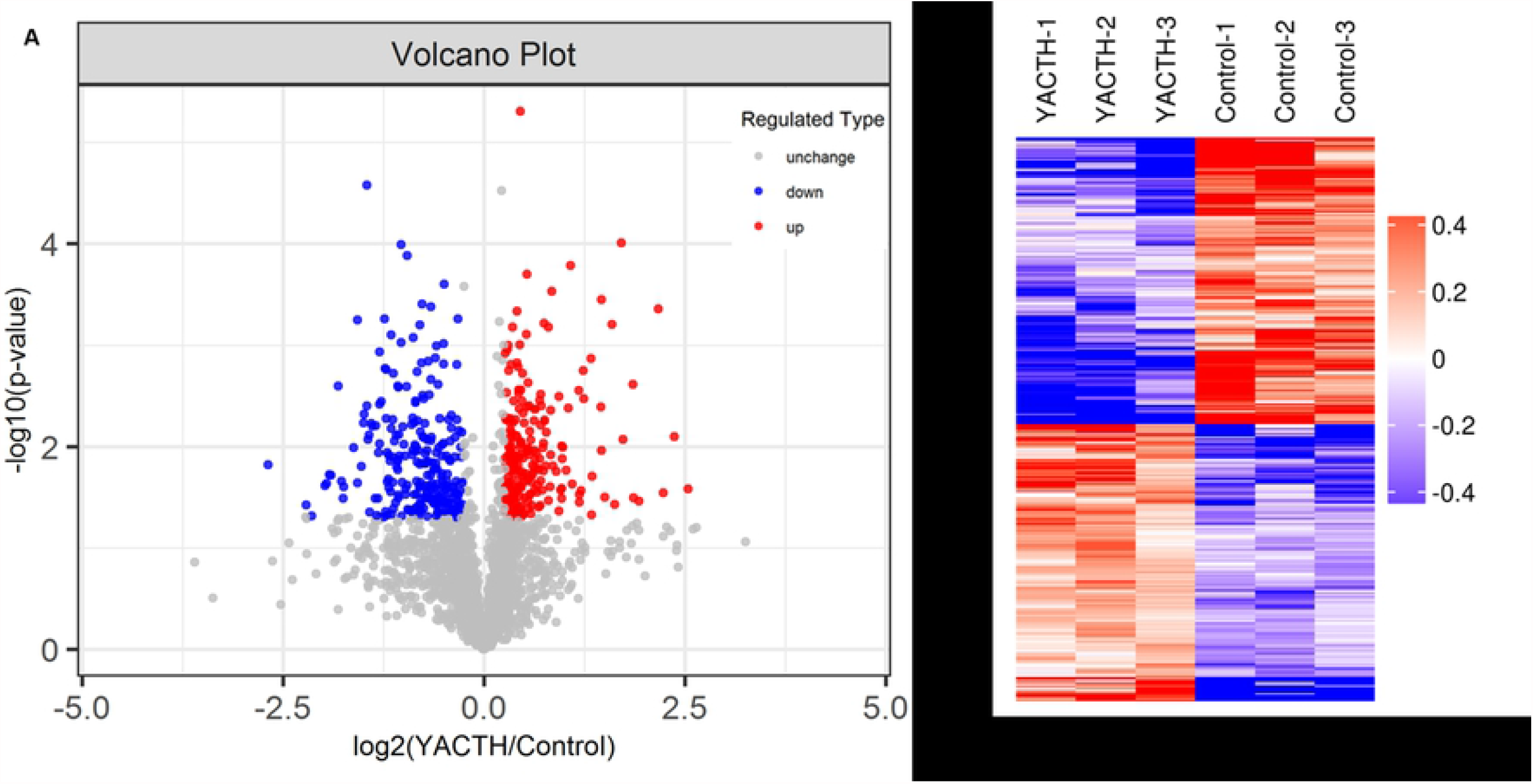
Volcano plot and heat map of SAU protein expressions under YACTH treatment.

Fig 2A is the volcano plot of the proteins that were affected by the YACTH treatment. While the grey dots represent the proteins that remained unchanged, the red dots represent the upregulated proteins and the blue dots represent the down-regulated proteins in the presence of YACTH. Fig 2B is the heat map that depicts the proteins being influenced by YACTH treatment. Each column represents a group of proteins. The protein expression levels are normalized by log2 and labeled in different colors. The up-regulated proteins are colored in red, and down-regulated proteins are colored in blue.

### Functional Analysis

In sub-cellular localization analysis, 384 cytoplasmic proteins, 83 membrane proteins, and 87 extracellular proteins were identified (S1 Fig). In addition, we conducted domain analysis to match YACTH regulated proteins with acknowledged structural domains. The majority of structural domains could be found in the proteins affected by YACTH treatment (Fig 3A). Typical structural domains such as ABC transporter, LPXTG cell wall anchor motif, YSIRK type signal peptide, radical SAM superfamily, and B domain, etc were observed. To study the overall response of SAU in the presence of YACTH, we performed GO analysis to assign the individual protein expression changes into certain groups. Take the example of the Top 20 enriched Go terms, proteins with significant expression changes were grouped into six biological processes with the highest number of proteins belongs to the cellular amide metabolic process, eight molecular functions with the highest number of proteins belong to RNA binding, six cellular components with the number of proteins almost evenly distributed in each cellular component (Fig 3B).

**Fig 3.**
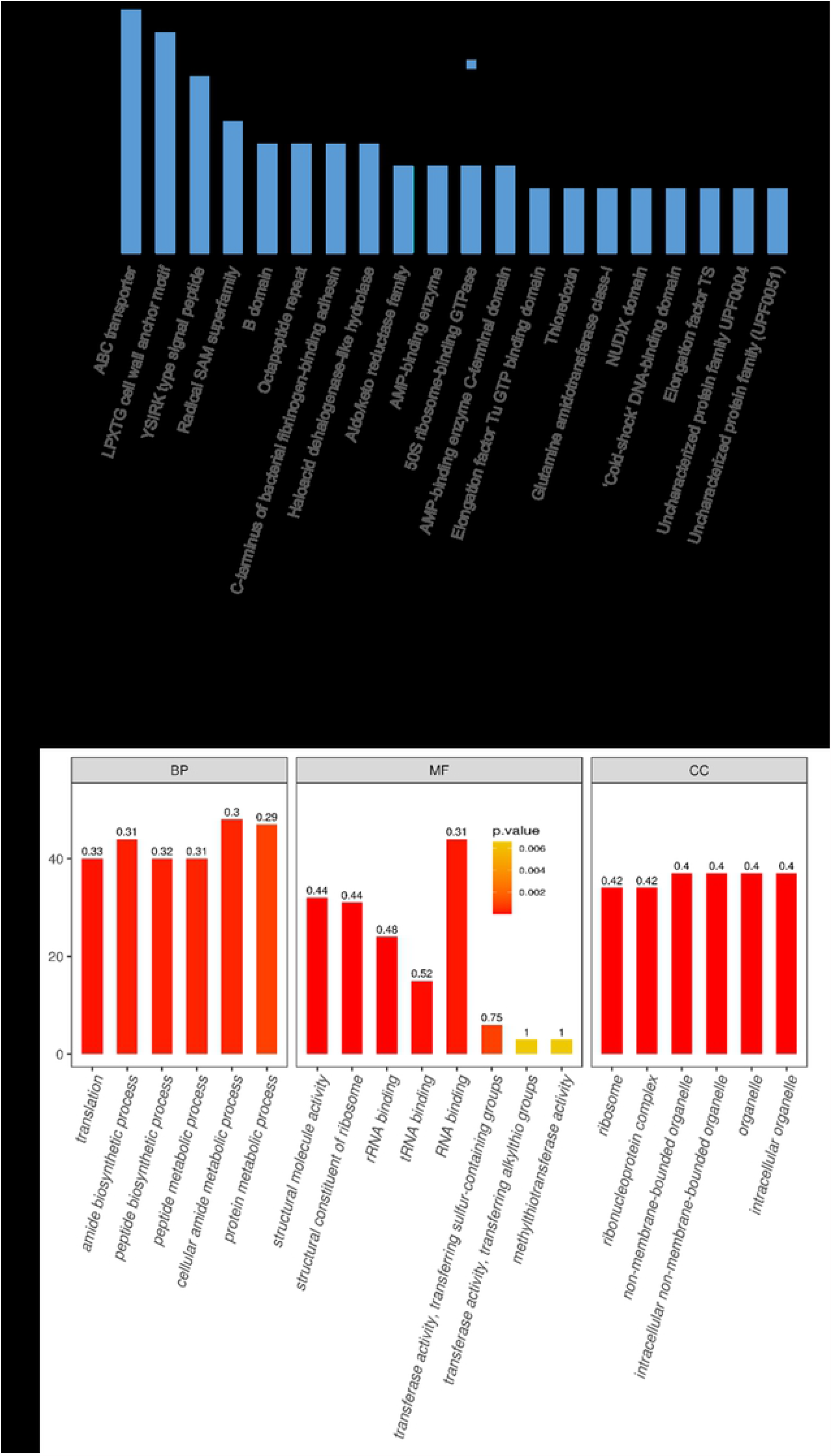
Domain analysis and GO analysis of the SAU proteins that were affected by the YACTH treatment.

Fig 3A shows the top 20 proteins that were matched with structural domains from the protein data bank. The length of the blue bars represents the number of proteins. Fig 3B depicts the GO enrichment analysis of the top 20 proteins. The proteins were categorized by biological process (BP), molecular function (MF), and cellular component (CC). The p-value is represented by the color intensity. The more intense red color represents a lower p-value.

In addition, we performed KEGG pathway analysis, which indicated that YACTH treatment regulated the expression of abundant SAU proteins, which were widely involved in a variety of biological processes and multiple metabolic pathways (Fig 4). The size of the dots represents the number of changed proteins, and the biggest protein number belongs to the ribosome, which is the key organelle responsible for translational activities. Other vital pathways related to SAU growth and metabolism, such as RNA polymerase, aminoacyl-tRNA biosynthesis, pyruvate metabolism, and aminoacyl-tRNA biosynthesis, were also presented in our data (Fig 4).

**Fig 4.**
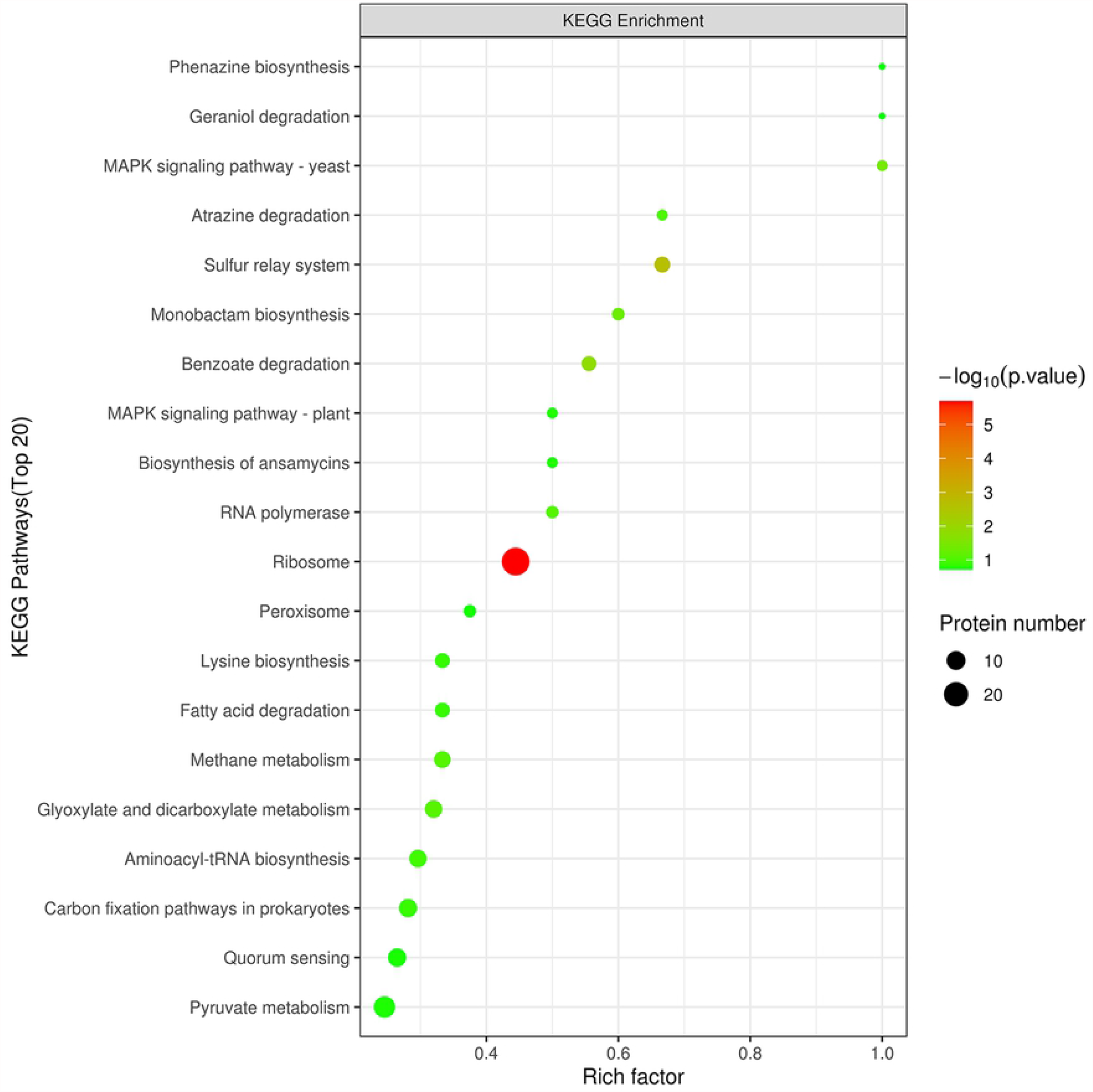
KEGG Pathway analysis of the SAU proteins that were affected by the YACTH treatment.

The proteins influenced by the YACTH treatment are assigned to different pathways, while the top 20 pathways are exhibited in Fig 4. Five grades (from 1 to 5) of the p-value, converted by –log10, were presented in colors while the size of the dots represents the number of proteins.

### PRM analysis

According to the GO and KEGG analysis of TMT results, 26 target proteins were selected to conduct PRM analysis; 6 of these proteins were significantly down-regulated and 20 proteins up-regulated as shown in Table 1. The up-regulated and down-regulated trends of all proteins of TMT were consistent with that of PRM analysis. For example, the DEAD-box RNA helicase cshA related to RNA degradation was up-regulated by 1.25-fold and 1.35-fold, while Putative DNA-directed RNA polymerase subunit delta related to RNA polymerase was down-regulated to 0.36 times and 0.07 times in TMT and PRM analysis, respectively. The up-regulated proteins were primarily involved in the pathways related to the ribosome, mismatch repair, aminoacyl-tRNA biosynthesis, and ABC transporter, etc. Three proteins associated with Mismatch repair were up-regulated 1.25-1.27 times. Six proteins in the ribosome, two proteins in ABC transporter, and one protein in glycolysis were also up-regulated. On other hand, the down-regulated proteins were mainly involved in the pathways related to RNA polymerase and the two-component system. At the same time, some pathways were regulated in both up-and down-directions. In glycolysis, three proteins were upregulated, while two proteins were down-regulated; in ABC transporter, two proteins were up-regulated, while three proteins were down-regulated. All the TMT results are statistically significant, and PRM analysis further confirmed the results of TMT (Table 1).

**Table 1.** PRM analysis of SAU proteins that were affected by the YACTH treatment.

| Map Name | Protein Name |  | Gene Name | TMT |  | PRM |
| --- | --- | --- | --- | --- | --- | --- |
|  |  |  |  | Ratio<br>YACTH/Control | TTEST<br>YACTH/Control | Ratio<br>YACTH/Control |
| Mismatch repair | A0A0E0VPG2 | RecJ | ST398NM01_1699 | 1.27 | 0.027 | 1.46 |
|  | Q6GFF3 | DNA ligase | ligA | 1.25 | 0.025 | 1.29 |
|  | P64319 | ATP-dependent DNA helicase PcrA | pcrA | 1.25 | 0.01 | 1.33 |
| RNA degradation | A0A077UMU7 | DEAD-box ATP-dependent RNA helicase CshA | cshA | 1.25 | 0.049 | 1.35 |
| RNA polymerase | P66725 | DNA-directed RNA polymerase subunit omega | rpoZ | 0.67 | 0.045 | 0.23 |
|  | A0A5S9C4T9 | Putative DNA-directed RNA polymerase subunit delta | rpoE | 0.36 | 0.004 | 0.07 |
| Ribosome | A7X5D9 | 50S ribosomal protein L18 | rplR | 1.81 | 0.016 | 1.94 |
|  | A0A033UZC9 | 30S ribosomal protein S21 | rpsU | 1.6 | 0.017 | 2.04 |
|  | A0A2D1CK30 | 30S ribosomal protein S10 | rpsJ | 1.45 | 0.001 | 1.57 |
|  | A0A077VMB7 | 50S ribosomal protein L21 | rplU | 1.42 | 0.032 | 1.38 |
|  | Q2YYQ1 | 50S ribosomal protein L22 | rplV | 1.39 | 0.009 | 2.57 |
|  | A0A0H2HWH9 | 50S ribosomal protein L14 | rplN | 1.24 | 0.036 | 1.83 |
|  | Q6GJC3 | 30S ribosomal protein S12 | rpsL | 0.81 | 0.041 | 0.45 |
| Glycolysis | A0A0E0VRM3 | Acetyl-coenzyme A synthetase | ST398NM01_1785 | 0.62 | 0.001 | 0.23 |
| Oxidative phosphorylation | A0A0E1XGX4 | Succinate dehydrogenase iron-sulfur subunit | sdhB | 0.82 | 0.013 | 0.69 |
| ABC transporters | A0A454GWW4 | SAV1866 family putative multidrug efflux ABC transporter (Fragment) | C7P97_12990 | 1.26 | 0.012 | 3.09 |
|  | A0A0E1XEH6 | Periplasmic binding protein | HMPREF0769_10977 | 1.25 | 0.049 | 1.71 |
|  | A0A0E1XHE1 | ABC transporter, ATP-binding protein | HMPREF0769_11391 | 0.59 | 0 | 0.18 |
|  | A0A2X2K196 | Response regulator ArlR | arlR_1 | 0.66 | 0.001 | 0.56 |
|  | Q6GFH6 | Low molecular weight protein-tyrosine-phosphatase PtpA | ptpA | 0.58 | 0.025 | 0.31 |
|  | A0A1Q8DED0 | D-alanyl carrier protein | dltC | 0.44 | 0.047 | 0.06 |

Table 1 shows the 26 affected proteins related to 7 biological processes & metabolic pathways. All the TMT results are statistically significant( $p < 0.05$ ) and are confirmed by
PRM analysis. The ratio of down-regulated proteins in YACTH group compared to the control group is highlighted in green.

## Discussion

SAU is a gram-positive bacterium that causes many diseases [18], the current therapeutic strategies against SAU mainly target the cell envelope, protein synthesis, and nucleic acids biosynthesis of the bacteria [19]. With the extensive use of antibiotics, the emergence of drug-resistant strains has become a great challenge for their treatment, which urges people to explore new antibiotics [20]. The herbal formular of YACTH is widely used in Southern China to prevent post-traumatic infection which is commonly caused by SAU [21,22]. In the present study, proteomic analysis was used to analyze the effects of YACTH on the protein expression of SAU. The expressions of 490 proteins were significantly changed in YACTH treated group compared to the control group. These altered proteins were widely associated with a variety of biological processes and multiple metabolic pathways. Interestingly, the down-regulated proteins were mainly involved in the pathways related to RNA polymerase (RNAP), oxidative phosphorylation, and two-component system. The RNAP multisubunit complex is responsible for bacterial transcription [23]. It contains α (two copies), β, β’ and ω subunit (Fig 5). The small ω subunit, namely rpoZ, plays key roles in subunit folding, complex assembly, and maintenance of transcriptional integrity [24]. The δ factor, rpoE, plays an essential role in RNA polymerase function in SAU [25]. The ablation of rpoE resulted in decreased transcription, protein synthesis, and pathogenicity [26]. The decreased expression of rpoZ and rpoE in SAU exposed to YACTH suggested that it may inhibit SAU’s activities on transcription level by RNAP pathway. In addition, YACTH significantly increased the level of the DEAD-box RNA helicase cshA, which was responsible for RNA degradation and might regulate the expression of virulence factors [27].

**Figure 5.**
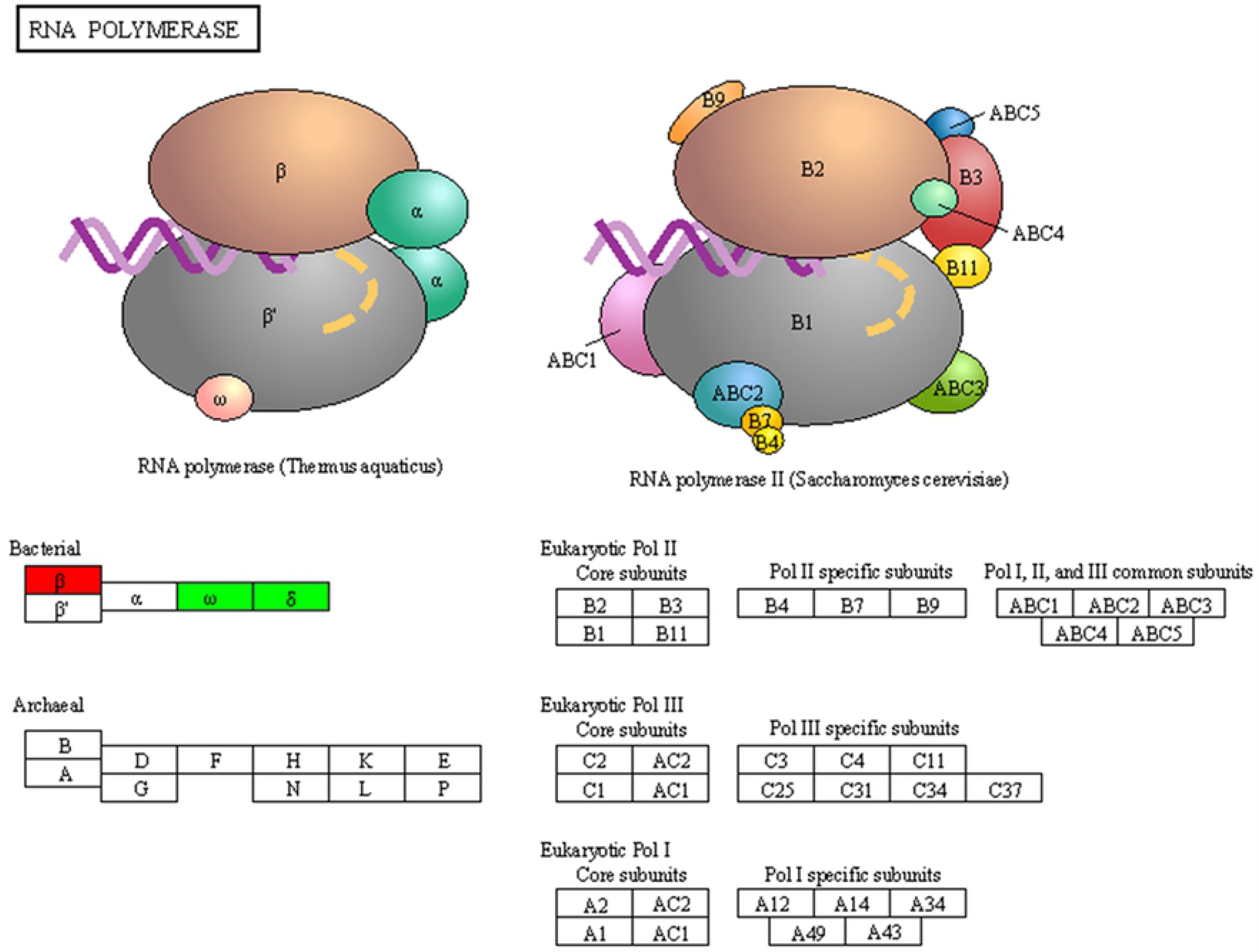
RNA polymerase of bacteria and eukaryote.

YACTH significantly down-regulated the expression of several proteins in two-component system and oxidative phosphorylation pathways, including arlRs, ptpA, dltC, and sdhB, etc (Table1), which were potential antibiotics targets associated with bacterial virulence and biofilm formation of SAU [28,29]. The two-component system signal transduction plays a pivotal role in regulating virulence gene expression, cell wall synthesis, biofilm formation, by which bacteria adapt to the external environment [30]. The biofilm formation of SAU is closely related to the intercellular communication and self-defense of the bacteria against antibiotics [31]. For example, arlRs is an antibiotic target for oxacillin, deletion of the arlRs in SAU increased its susceptibilities to oxacillin [32]. The protein tyrosine phosphatase ptpA is a signaling molecule and may interfere with host cell signaling [29,33], which contributes to the infectivity of SAU [34]. Deletion of ptpA not only impaired the survival rates of SAU in macrophages but also reduce the SAU infectivity on mice [29]. DltC, a two-component system protein that affects the cell surface charge of bacteria, is involved in the D-alanine substitution of lipoteichoic acid, and therefore biofilm formation [35]. dltC regulates D-alanine incorporation into cell wall polymers in SAU. A lack of this decoration leads to increased susceptibility to cationic antimicrobial peptide [36,37]. Another downregulated protein associated with biofilm formation is sdhB, a succinate dehydrogenase iron-sulfur subunit involved in the oxidation phosphorylation pathway[38]. The sdhB gene knockout resulted in a 1.2 fold significant decrease of biofilm formation[39], while mutation of sdhB led to defect in the persistence of levofoxacin[38].

On other hand, the up-regulated proteins were primarily involved in the pathways related to mismatch repair, ribosome, aminoacyl-tRNA biosynthesis, and ABC transporter (Table 1). The up-regulated mismatch repair proteins included pcrA, recJ, and ligA. ATP-dependent DNA helicase pcrA is an essential helicase in Gram-positive bacteria, genetic induction experiment indicated that overproduction of pcrA has deleterious effects on the host cell and plasmid replication [40]. Single-stranded-DNA-specific exonuclease recJ is involved in a variety of DNA repair pathways corresponding to their cleavage polarities, but the relationship between the cleavage polarity and the respective DNA repair pathways has not been clarified [41]. NAD-dependent DNA ligase (ligA) has been thought of as a potential broad-spectrum antibacterial target [42,43,44] and was later proved to be an antibacterial target of SAU and inhibited by pyridochromanone[42]. Our result suggested that YACTH impaired SAU and therefore led to stress response with increased expression of proteins that are involved in mismatch repair. For ribosomes, eighteen ribosomal proteins were upregulated, while only one ribosomal protein was downregulated in YACTH treatment group (table1). Among the 18 upregulated ribosomal proteins, four are related to antibiotics sensitivity, as was reported in previous publications. Mutations of 30S ribosomal proteins S21, rpsU, led to reduced susceptivity of vancomycin of SAU [45]. Increased level of rpIN was also observed in vancomycin-intermediate SAU (VISA) [46]. Mutations on 30S ribosomal proteins S10, rpsJ, was reported to be the cause of reduced tigecycline susceptivity [47]. While mutation of 50S ribosomal protein L22, rplV, is related to several antibiotic resistance, high-level telithromycin selection led to the mutation on rplV [48], mutations of rplV reduced sensitivity of SAU to erythromycin and other macrolides were not only observed in the laboratory but also clinical studies [49,50,51].

As mentioned above, we reported a new antimicrobial herbal formularlation of YACTH and its effects on the expression of bacterial proteins, which might be an alternative choice to protect against post-traumatic infection caused by SAU. The multiple proteins down-regulated by YACTH were primarily involved in the RNA polymerase, oxidative phosphorylation, and two-component system pathways, which are associated with the biofilm formation and the virulence secretion of SAU. Otherwise, up-regulated proteins by YACTH were mainly presented in the pathways of mismatch repair, ribosome aminoacyl-tRNA biosynthesis, and ABC transporter, etc. Some down- and up-regulated proteins were found to be related to antibiotic sensitivity. These results provided pieces of evidence for the hypothesis that YACTH might repress the growth and infectivity of SAU by either reducing or increasing the expression of target proteins. Some of these proteins were associated with cell envelope, protein synthesis, and nucleic acid biosynthesis, which are pathways that are targets of traditional antibiotics. Unlike common antibiotics that solely mediated on one specific target, YACTH regulated multiple proteins that are involved in several signaling pathways, probably due to its complex components. Such results provided a broad direction of exploring new anti-bacterial targets to fight against SAU. Further studies were needed to clarify the preliminary antibacterial mechanism of each effective component, structure, and potential targets.

## Supporting Information

**S1 Fig. Subcellular localization of the SAU proteins that were affected by the YACTH treatment**.

The SAU proteins that were affected by YACTH treatment were assigned into three subcellular locations: cytoplasmic(384), extracellular(87), and membrane(83).

## Reference

1. Wertheim HF, Melles DC, Vos MC, van Leeuwen W, van Belkum A, Verbrugh HA, et al. The role of nasal carriage in Staphylococcus aureus infections. Lancet Infect Dis. 2005;5(12):751–62. doi: 10.1016/S1473-3099(05)70295-4.

2. Sabike II, Fujikawa H, Sakha MZ, Edris AM. Production of Staphylococcus aureus enterotoxin a in raw milk at high temperatures. J Food Prot. 2014;77(9):1612–6. doi: 10.4315/0362-028X.JFP-13-527.

3. Aggarwal S, Jena S, Panda S, Sharma S, Dhawan B, Nath G, et al. Antibiotic susceptibility, virulence pattern, and typing of Staphylococcus aureus strains Isolated from variety of infections in India. Front Microbiol. 2019;10:2763. doi: 10.3389/fmicb.2019.02763.

4. Nowicka D, Grywalska E. Staphylococcus aureus and host immunity in recurrent furunculosis. Dermatology. 2019;235(4):295–305. doi: 10.1159/000499184. Epub 2019 Apr 17.

5. Yang L, Feng J, Liu J, Yu L, Zhao C, Ren Y,et al. Pathogen identification in 84 Patients with post-traumatic osteomyelitis after limb fractures. Ann Palliat Med 2020;9(2):451–458. doi: 10.21037/apm.2020.03.29

6. Li Z, Zheng F, C Y, Li H, Liu D, LiuX. Optimization of the preperation process of Shangkehuangshui II.ChinHospPharmJ, 2018;38(24):2532-36.doi:10.13286/J.cnki.chinhosppharmacyj.2018.24.07

7. Xu H, Li Z, Lei K, Li H, Liu D, Chen Y, et al. Study on the effect of Shangkehuangshui on acute soft tissue injury. Pharmacology and clinic of traditional Chinese medicine 2017; 33 (6): 124–130

8. Cai L, Yang H, Sun B, Wen J, WuF, Wu F, Y J. Effect of the traumatology yellow water on new bone formation of distraction osteogenesis zone in rabbit. Bone Setting of Traditional Chinese medicine. 2014; 26 (10): 12–15.

9. Kälicke T, Schlegel U, Printzen G, Schneider E, Muhr G, Arens S. Influence of a standardized closed soft tissue trauma on resistance to local infection. An experimental study in rats. J Orthop Res. 2003, 21(2):373–8. doi: 10.1016/S0736-0266(02)00149

10. Giesbrecht P, Kersten T, Maidhof H et al. Staphylococcal cell wall:morphogenesis and fatal variations in the presence of penicillin. Microbiol Mol Biol R 1998;62:1371–41

11. Nguyen F, Starosta AL, Arenz S et al. Tetracycline antibiotics and resistance mechanisms. Biol Chem 2014;395:p559–75.

12. Foster TJ. Antibiotic resistance in Staphylococcus aureus. Current status and future prospects. FEMS Microbiol Rev. 2017 May 1;41(3):430–449.

13. Schulze WX, Usadel B. Quantitation in mass-spectrometry-based proteomics. Annu Rev Plant Biol. 2010;61:p491–516.

14. Aebersold R, Burlingame AL, Bradshaw RA. Western blots versus selected reaction monitoring assays: time to turn the tables? Mol Cell Proteomics. 2013 Sep;12(9):2381–2. doi: 10.1074/mcp.E113.031658.

15. Cao J, Fu H, Gao L, Zheng Y. Antibacterial activity and mechanism of lactobionic acid against Staphylococcus aureus. Folia Microbiol (Praha). 2019 Nov;64(6):899–906

16. Jeong YI, Na HS, Seo DH, Kim DG, Lee HC, Jang MK, et al. Ciprofloxacin-encapsulated poly(DL-lactide-co-glycolide) nanoparticles and its antibacterial activity. Int J Pharm. 2008 Mar 20;352(1-2):317–23. doi: 10.1016/j.ijpharm.2007.11.001.

17. CLSI Guidelines (2018) Performance Standards for Antimicrobial Susceptibility Testing. 28th Edition (M100S).

18. Oliveira D, Borges A, Simões M. Staphylococcus aureus Toxins and Their Molecular Activity in Infectious Diseases. Toxins (Basel). 2018 Jun 19;10(6):252. doi: 10.3390/toxins10060252.

19. Assis LM, Nedeljković M, Dessen A. New strategies for targeting and treatment of multi-drug resistant Staphylococcus aureus. Drug Resist Updat. 2017 Mar;31:1–14. doi: 10.1016/j.drup.2017.03.001.

20. Brown ED, Wright GD. Antibacterial drug discovery in the resistance era. Nature. 2016 Jan 21;529(7586):336–43. doi: 10.1038/nature17042. PMID: 26791724.

21. Helmann JD. RNA polymerase: a nexus of gene regulation. Methods. 2009 Jan;47(1):1–5. doi: 10.1016/j.ymeth.2008.12.001. PMID: 19070783; PMCID: PMC3022018.

22. Weiss A, Shaw LN. Small things considered: the small accessory subunits of RNA polymerase in Gram-positive bacteria. FEMS Microbiol Rev. 2015 Jul;39(4):541–54. doi: 10.1093/femsre/fuv005.

23. Mathew R, Chatterji D. The evolving story of the omega subunit of bacterial RNA polymerase. Trends Microbiol. 2006 Oct;14(10):450–5. doi: 10.1016/j.tim.2006.08.002.

24. Weiss A, Moore BD, Tremblay MHJ, Chaput D, Kremer A, Shaw LN. The ω subunit governs RNA polymerase stability and transcriptional specificity in Staphylococcus aureus. J Bacteriol. 2016 Dec 28;199(2):e00459–16. doi: 10.1128/JB.00459-16. PMID: 27799328; PMCID: PMC5198492.

25. Lin Z, Wang F, Shang Z, Lin W. Biochemical and structural analyses reveal critical residues in δ subunit affecting its bindings to β’ subunit of Staphylococcus aureus RNA polymerase. Biochem Biophys Res Commun. 2021 Mar 19;545:98–104. doi: 10.1016/j.bbrc.2021.01.078.

26. Weiss A, Ibarra JA, Paoletti J, Carroll RK, Shaw LN. The δ subunit of RNA polymerase guides promoter selectivity and virulence in Staphylococcus aureus. Infect Immun. 2014 Apr;82(4):1424–35. doi: 10.1128/IAI.01508-14.

27. Oun S, Redder P, Didier JP, François P, Corvaglia AR, Buttazzoni E, Giraud C, Girard M, Schrenzel J, Linder P. The CshA DEAD-box RNA helicase is important for quorum sensing control in Staphylococcus aureus. RNA Biol. 2013 Jan;10(1):157–65. doi: 10.4161/rna.22899.

28. Horn J, Stelzner K, Rudel T, Fraunholz M. Inside job: Staphylococcus aureus host-pathogen interactions. Int J Med Microbiol. 2018 Aug;308(6):607–624. doi: 10.1016/j.ijmm.2017.11.009. Epub 2017 Nov 26. PMID: 29217333.

29. Gannoun-Zaki L, Pätzold L, Huc-Brandt S, Baronian G, Elhawy MI, Gaupp R, et al. PtpA, a secreted tyrosine phosphatase from Staphylococcus aureus, contributes to virulence and interacts with coronin-1A during infection. J Biol Chem. 2018 Oct 5;293(40):15569–15580. doi:10.1074/jbc.RA118.003555.

30. Kuroda M, Ohta T, Uchiyama I, Baba T, Yuzawa H, Kobayashi I, et al. Whole genome sequencing of meticillin-resistant Staphylococcus aureus. Lancet. 2001 Apr 21;357(9264):1225–40. doi: 10.1016/s0140-6736(00)04403-2.

31. Schilcher K, Horswill AR. Staphylococcal biofilm development: structure, regulation, and treatment strategies. Microbiol Mol Biol Rev. 2020 Aug 12;84(3):e00026–19. doi: 10.1128/MMBR.00026-19.

32. Bai J, Zhu X, Zhao K, Yan Y, Xu T, Wang J, et al. The role of ArlRS in regulating oxacillin susceptibility in methicillin-resistant Staphylococcus aureus indicates it is a potential target for antimicrobial resistance breakers. Emerg Microbes Infect. 2019;8(1):503–515. doi: 10.1080/22221751.2019.1595984.

33. Brelle S, Baronian G, Huc-Brandt S, Zaki LG, Cohen-Gonsaud M, Bischoff M, et al. Phosphorylation-mediated regulation of the Staphylococcus aureus secreted tyrosine phosphatase PtpA. Biochem Biophys Res Commun. 2016 Jan 15;469(3):619–25. doi: 10.1016/j.bbrc.2015.11.123. Epub 2015 Dec 8.

34. Vega C, Chou S, Engel K, Harrell ME, Rajagopal L, Grundner C. Structure and substrate recognition of the Staphylococcus aureus protein tyrosine phosphatase PtpA. J Mol Biol. 2011 Oct 14;413(1):24–31. doi: 10.1016/j.jmb.2011.08.015.

35. Matsuo M, Oogai Y, Kato F, Sugai M, Komatsuzawa H. Growth-phase dependence of susceptibility to antimicrobial peptides in Staphylococcus aureus. Microbiology (Reading). 2011 Jun;157(Pt 6):1786–1797. doi: 10.1099/mic.0.044727-0.

36. Reichmann NT, Cassona CP, Gründling A. Revised mechanism of D-alanine incorporation into cell wall polymers in Gram-positive bacteria. Microbiology (Reading). 2013 Sep;159(Pt 9):1868–1877. doi: 10.1099/mic.0.069898-0.

37. Wood BM, Santa Maria JP Jr, Matano LM, Vickery CR, Walker S. A partial reconstitution implicates DltD in catalyzing lipoteichoic acid d-alanylation. J Biol Chem. 2018 Nov 16;293(46):17985–17996. doi: 10.1074/jbc.RA118.004561.

38. Wang W, Chen J, Chen G, Du X, Cui P, Wu J, et al. Transposon mutagenesis identifies novel genes associated with Staphylococcus aureus persister formation. Front Microbiol. 2015 Dec 23;6:1437. doi: 10.3389/fmicb.2015.01437.

39. De Backer S, Sabirova J, De Pauw I, De Greve H, Hernalsteens JP, Goossens H,et al. Enzymes catalyzing the tca-and urea cycle influence the matrix composition of biofilms formed by methicillin-resistant Staphylococcus aureus USA300. Microorganisms. 2018 Oct 29;6(4):113. doi: 10.3390/microorganisms6040113.

40. Iordanescu S. Characterization of the Staphylococcus aureus chromosomal gene pcrA, identified by mutations affecting plasmid pT181 replication. Mol Gen Genet. 1993 Oct;241(1-2):185–92. doi: 10.1007/BF00280216.

41. Shimada A, Masui R, Nakagawa N, Takahata Y, Kim K, Kuramitsu S, et al. A novel single-stranded DNA-specific 3’-5’ exonuclease, Thermus thermophilus exonuclease I, is involved in several DNA repair pathways. Nucleic Acids Res. 2010 Sep;38(17):5692–705. doi: 10.1093/nar/gkq350.

42. Dwivedi N, Dube D, Pandey J, Singh B, Kukshal V, Ramachandran R, et al. NAD(+)-dependent DNA ligase: a novel target waiting for the right inhibitor. Med Res Rev. 2008 Jul;28(4):545–68. doi: 10.1002/med.20114.

43. Podos SD, Thanassi JA, Pucci MJ. Mechanistic assessment of DNA ligase as an antibacterial target in Staphylococcus aureus. Antimicrob Agents Chemother. 2012 Aug;56(8):4095–102. doi: 10.1128/AAC.00215-12.

44. Kaczmarek FS, Zaniewski RP, Gootz TD, Danley DE, Mansour MN, Griffor M, et al. Cloning and functional characterization of an NAD(+)-dependent DNA ligase from Staphylococcus aureus. J Bacteriol. 2001 May;183(10):3016–24. doi: 10.1128/JB.183.10.3016-3024.2001.

45. Blake KL, O’Neill AJ. Transposon library screening for identification of genetic loci participating in intrinsic susceptibility and acquired resistance to antistaphylococcal agents. J Antimicrob Chemother. 2013 Jan;68(1):12–6. doi: 10.1093/jac/dks373. Epub 2012 Oct 7.

46. Sirichoat A, Lulitanond A, Kanlaya R, Tavichakorntrakool R, Chanawong A, Wongthong S, et al. Phenotypic characteristics and comparative proteomics of Staphylococcus aureus strains with different vancomycin-resistance levels. Diagn Microbiol Infect Dis. 2016 Dec;86(4):340–344. doi: 10.1016/j.diagmicrobio.2016.09.011.

47. Beabout K, Hammerstrom TG, Perez AM, Magalhães BF, Prater AG, Clements TP, et al. The ribosomal S10 protein is a general target for decreased tigecycline susceptibility. Antimicrob Agents Chemother. 2015 Sep;59(9):5561–6. doi: 10.1128/AAC.00547-15.

48. Gentry DR, Holmes DJ. Selection for high-level telithromycin resistance in Staphylococcus aureus yields mutants resulting from an rplB-to-rplV gene conversion-like event. Antimicrob Agents Chemother. 2008 Mar;52(3):1156–8. doi: 10.1128/AAC.00923-07. Epub 2008 Jan 14.

49. Prunier AL, Malbruny B, Laurans M, Brouard J, Duhamel JF, Leclercq R. High rate of macrolide resistance in Staphylococcus aureus strains from patients with cystic fibrosis reveals high proportions of hypermutable strains. J Infect Dis. 2003 Jun 1;187(11):1709–16. doi: 10.1086/374937.

50. Prunier AL, Malbruny B, Tandé D, Picard B, Leclercq R. Clinical isolates of Staphylococcus aureus with ribosomal mutations conferring resistance to macrolides. Antimicrob Agents Chemother. 2002 Sep;46(9):3054–6. doi: 10.1128/aac.46.9.3054-3056.2002.

51. Han D, Liu Y, Li J, Liu C, Gao Y, Feng J, et al. Twenty-seven-nucleotide repeat insertion in the rplV gene confers specific resistance to macrolide antibiotics in Staphylococcus aureus. Oncotarget. 2018 May 25;9(40):26086–26095. doi: 10.18632/oncotarget.25441.

